# Chitosan inhibits septin-mediated plant infection by the rice blast fungus *Magnaporthe oryzae* in a Protein Kinase C and Nox1 NADPH oxidase-dependent manner

**DOI:** 10.1101/2020.05.15.098657

**Authors:** Federico Lopez-Moya, Magdalena Martin-Urdiroz, Miriam Oses-Ruiz, Mark D. Fricker, George R. Littlejohn, Luis V. Lopez-Llorca, Nicholas J. Talbot

## Abstract

- Chitosan is a partially deacetylated linear polysaccharide composed of β-1,4-linked units of D-glucosamine and N-acetyl glucosamine. As well as acting as a structural component of fungal cell walls, chitosan can be applied as a potent antifungal agent. However, the mode-of-action of chitosan in fungal pathogens is poorly understood.
- Here, we report that chitosan is effective for control of rice blast disease. Chitosan application impairs growth of the blast fungus *Magnaporthe oryzae* and has a pronounced effect on appressorium-mediated plant infection. Chitosan inhibits septin-mediated F-actin re-modelling at the appressorium pore, thereby preventing re-polarisation of the infection cell and rice leaf cuticle penetration.
- We found that chitosan causes plasma membrane permeabilization of *M. oryzae* and affects NADPH oxidase-dependent synthesis of reactive oxygen species, essential for septin ring formation and fungal pathogenicity. Our data further show that the toxicity of chitosan to *M. oryzae* requires the protein kinase C-dependent cell wall integrity pathway and the Nox1 NADPH oxidase. A conditionally lethal, analogue (PP1)-sensitive mutant of Pkc1 is partially remediated for growth in the presence of chitosan and PP1, while Δ*nox1* mutants increase their glucan/chitin cell wall ratio, rendering them resistant to chitosan.
- Taken together, our data show that chitosan is a potent fungicide for control of the rice blast fungus which involves the cell wall integrity pathway, disrupts plasma membrane and inhibits septin-mediated plant infection.

## Introduction

Plant pathogenic fungi are responsible for many of the world’s most serious crop diseases and yet remain very challenging to control (Fisher *et al.*, 2012; Fisher *et al*. 2018). Fungicide application is often not completely effective, has environmental consequences and resistance to fungicides occurs very frequently (Lucas *et al.*, 2015). Chitosan is a biopolymer obtained by partial N-deacetylation of the β-1,4-D-linked polymer of N-acetyl glucosamine, chitin (Kumar, 2000). As well as being a structural component of fungal cell walls, chitosan also displays anti-microbial activity (Allan & Hadwiger, 1979). Chitosan has a pK_a_ value of ∼6.3 and it is cationic at lower pH values, due to protonation of its amino groups. Chitosan inhibits the growth of filamentous fungal plant pathogens, such as *Botrytis cinerea* (Muñoz & Moret, 2010) and *Fusarium oxysporum* (Palma-Guerrero *et al.*, 2008; Al-Hetar *et al.*, 2010). Chitosan therefore shows considerable potential as a naturally occurring, novel anti-fungal agent, but its precise mode of action in plant pathogenic fungi remains unclear.

Chitosan inhibits the growth of sensitive fungi, causing massive membrane permeabilization (Palma-Guerrero *et al.*, 2009; Palma-Guerrero *et al.*, 2010). In *Neurospora crassa*, chitosan exposure causes generation of an oxidative response, which leads to membrane permeabilization and cell death (Lopez-Moya *et al.*, 2015). Recent studies, however, have also demonstrated that cell wall composition plays a key role in fungal sensitivity to chitosan (Aranda-Martinez *et al.*, 2016) as well as mitochondrial activity (Jaime *et al.*, 2012). Transcriptional profiling studies meanwhile have revealed that cytoskeletal functions are also compromised by chitosan exposure in *N. crassa.* Genes related to actin polymerisation in *N. crassa*, for example, are repressed by chitosan treatment (Lopez-Moya, *et al.*, 2016), suggesting that actin-dependent functions, such as cell polarity, exocytosis, endocytosis, cytokinesis, and organelle movement (Barja *et al.*, 1993, Berepiki *et al.*, 2010) may be affected by chitosan exposure. In all of these reports, however, it is unclear whether the observations reveal the mode-of-action of chitosan, or rather the pleiotropic effects it has on cell viability. It is therefore vital to carry out more comprehensive investigations of the manner in which chitosan affects fungal viability, in order to determine its likely efficacy as a novel fungicide.

In this report, we describe a series of experiments designed to determine the mode-of-action of chitosan in the control of a major crop disease-causing fungus, *Magnaporthe oryzae*, which has a global impact on food security (Castroagudin *et al.*, 2015; Zhang *et al.*, 2009; Wilson & Talbot, 2009). *Magnaporthe oryzae* is the causal agent of rice blast disease and is responsible for up to 30% losses to the annual rice harvest (Talbot, 2003). *M. oryzae* infects rice cells by using specialised infection structures called appressoria. These structures generate enormous turgor, which is applied as physical force to penetrate epidermal cells and then infect rice tissues (Wilson & Talbot, 2009). The dome-shaped appressorium possesses a thick melanin layer in its cell wall, which is critical for infection (Chumley & Valent, 1990) and can generate up to 8.0 MPa of pressure (Howard *et al.*, 1991; Howard & Valent, 1996), by accumulating molar concentrations of glycerol and other polyols as compatible solutes (de Jong *et al*., 1997). Appressorium turgor is sensed by the Sln1 histidine-aspartate kinase, which acts via the Protein kinase C-dependent cell wall integrity pathway and the cAMP-dependent protein kinase A pathway in order to modulate glycerol accumulation and melanin biosynthesis, once a critical threshold of pressure has been reached. An NADPH oxidase-dependent regulated burst of reactive oxygen species (ROS) then occurs, which leads to septin-dependent re-polarisation of the appressorium and plant infection (Ryder *et al*., 2019). ROS generation requires NADPH oxidases encoded by the *NOX1, NOX2* and *NOXR* genes, which are all essential for *M. oryzae* infection (Egan *et al.*, 2007; Ryder *et al.*, 2013). In other fungi, NADPH oxidase-dependent ROS generation is also known to play roles in cell polarity and invasive growth. In *Sordaria macrospora*, for instance, Nox1 regulates gene expression involved in cytoskeleton remodelling, hyphal fusion and mitochondrial respiration (Dirschnabel *et al.*, 2014), while in the endophytic fungus *Epichloë festucae NOXA*, is essential for polarized growth and hyphal fusion (Eaton *et al.*, 2011). In *M. oryzae, NOX1*, is also involved in cell wall organisation and *Δnox1* mutants is resistant to many cell wall perturbing agents, such as calcofluor white (Egan *et al.*, 2007). This role may be conserved in fungi, because in the mycoparasitic fungus *Coniothyrium minitans*, for example, *CmNOX1* and *CmSLT2*, regulate localisation of the cell wall integrity-associated MAP kinase (MAPK) and mediate changes in fungal gene expression associated with cell wall integrity (Wei *et al.*, 2016).

The Nox2-regulated synthesis of ROS is necessary for recruitment and organisation of four septin guanine nucleotide-binding proteins to the appressorium pore (Ryder *et al*., 2013), where they form a hetero-oligomeric ring complex, essential for re-polarisation of the appressorium (Dagdas *et al.* 2012). The septin ring rigidifies the cortex of the appressorium, acting as a scaffold for F-actin organisation. Septins also act as a lateral diffusion barrier, required for localisation of polarity determinants such as Chm1 (Li *et al.* 2004), Tea1 and Las17 (Dagdas *et al.*, 2012).

In this study, we set out to determine the effect of exposure to chitosan on infection-related development by *M. oryzae*. Chitosan is known to be a cell wall (CW) component in *M. oryzae*, which is important in adhesion and appressorium formation (Geoghegan & Gurr, 2016). We were interested in whether exposure to exogenous chitosan was, however, toxic to the fungus, as shown for other fungi and, if so, how it might be acting on the cellular events necessary for plant infection. We provide evidence that chitosan exposure is fungicidal to *M. oryzae.* causing membrane permeabilization and preventing plant infection. Importantly, we also show that the cell wall integrity pathway and Nox1 NADPH oxidase activity are both essential for the toxicity of chitosan, providing insight into its mode of action.

## Material and Methods

### Fungal strains and growth conditions

The wild-type strain of *Magnaporthe oryzae*, Guy11 (Leung *et al*., 1988) and transgenic rice blast isolates expressing Sep4-GFP::H1-RFP, Sep5-GFP, Gelsolin-GFP, Chm1-GFP, Tea1-GFP and Grx1-roGFP2 strains were stored in the laboratory of NJT. Gene replacement mutants Δ*nox1*, Δ*nox2*, Δ*nox1nox2*, Δ*mps1* and the *pkc1*^*AS*^ were generated as described previously (Xu *et al*., 1998; Ryder *et al*., 2013; Penn *et al*.,2015). All fungal strains were grown on complete medium (CM) at 24°C under a 12h light/dark photoperiod (Talbot *et al.*, 1993). Conidial suspensions were obtained in sterile distilled water (SDW) by scraping the surface of 12-day-old plate cultures with a spreader, before being filtered through Miracloth (Calbiochem, USA) and concentrated by centrifugation (13,000 rpm, 1 min).

### Preparation of Chitosan

Medium molecular weight chitosan (70 kDa) with an 85% deacetylation degree (T8) was provided by Marine BioProducts GmbH (Bremerhaven, Germany). Chitosan solutions were prepared as described by Palma-Guerrero *et al.* (2008). The resulting solution was dialysed against distilled water and autoclaved. Chitosan solutions were stored at 4 °C until used, and never stored for longer than 30 days.

### Exposure of *M. oryzae* to chitosan during plant infection

To evaluate the effect of chitosan on the pathogenicity of *M. oryzae*, leaf spot and spray infection assays were performed. Conidia of Guy11 were collected and suspensions adjusted to 10^5^ conidia ml^-1^. Leaf spot bioassays were performed by inoculating 20µl droplets of 1×10^5^ conidia ml^-1^ onto the adaxial surfaces of detached rice leaves of the blast-susceptible dwarf Indica variety of rice CO-39 (Talbot *et al.*, 1993) using 4 replicate leaves per treatment. Inoculated rice leaves were incubated in moist chambers and after 5d, the size of the resulting rice blast disease lesion recorded. Treatments included a control experiment (no chitosan) and conidial suspensions containing either 1 or 5 mg ml^-1^ chitosan, respectively. For leaf spray bioassays, CO-39 rice plants at the three-leaf stage (normally, 21-days-old) were sprayed with 5 ml of a suspension of 5×10^4^ conidia ml^-1^ using an artist’s airbrush (Badger Air-Brush Co, Franklin, IL, USA). Plants were also sprayed with chitosan only (no conidia) and all experiments repeated three times. Inoculated plants were placed in plastic bags for 2 days to achieve high humidity and symptoms scored after 5 days. Leaves were collected and rice blast lesions quantified.

### Leaf sheath bioassays

Leaf sheath fragments (2-3 cm long) were obtained from 21-day-old seedlings of rice cultivar CO-39. Leaf sections were inoculated with 50 µl of a conidial suspension of 1×10^5^ conidia ml^-1^ and incubated in moist chambers for 30h at 24°C. Transverse sections were cut with a razor blade and mounted in water. Micrographs were recorded using an IX-81 Olympus inverted microscope. After 30h, the frequency of *M. oryzae*-rice cell invasion was recorded (n=50 cells observed, in three replicate experiments).

### Evaluation of the effect of chitosan exposure on growth and development of *M. oryzae*

Conidia of Guy11 were incubated on hydrophobic glass coverslips in the presence of 0, 0.1, 0.5, 1, 2 and 5 mg ml^-1^ chitosan. The frequency of conidial germination was determined after 2h. The frequency of appressorium development was scored after 4, 6, 8, 16 and 24h.

The effect of chitosan on mycelium growth of *M. oryzae* was tested as follows. Guy 11, Δ*nox1*, Δ*nox2*, Δ*nox1nox2*, Δ*mps1* and *pkc1*^*AS*^ were grown on CM liquid medium for 48h on an orbital shaker (150 rpm) at 24°C with a 12/12h photoperiod. Mycelium was then transferred to CM medium in the presence/absence of chitosan (0.5 and 1 mg ml^-1^) and incubated for a further 48h. Mycelium was then collected and washed twice with SDW before being lyophilized and the dry weight recorded.

### Physiological and cellular response of *M. oryzae* to chitosan during appressorium differentiation

Reporter strains Sep4-GFP::H1-RFP, Sep5-GFP, Gelsolin-GFP, Chm1-GFP, Tea1-GFP and Grx1-roGFP2 were used to evaluate the effect of chitosan on cytoskeletal organisation, plasma membrane integrity and cellular oxidative status. A conidial suspension of each strain was incubated in the presence 5 mg ml^-1^ or absence of chitosan and SDW for 4h, 6h, 8h and 24h to visualise appressorium development. FM4–64 (N-(3-triethylammoniumpropyl-)-4-(6(4-(diethyl amino) phenyl) hexatrienyl) pyridinium dibromide) was used to evaluate the integrity of the plasma membrane in *M. oryzae* appressoria exposed to 5 mg ml^-1^ chitosan. Germlings were observed 24h after inoculation. A Grx1-roGFP2 strain (Samalova *et al.*, 2014) was used to detect changes in glutathione oxidation. This had a glutaredoxine (Grx) subunit to improve response kinetics to oxidation in order to evaluate the oxidative response of *M. oryzae* appressoria to either 1 or 5 mg ml^-1^ chitosan (Samalova *et al.*, 2014). Maximum fluorescence is related to the accumulation of oxidized glutathione a measurement to determine ROS.

For epifluorescence microscopy, an IX-81 Olympus inverted microscope connected to a CoolSNAP HQ2 camera was used. Three-dimensional projections were captured using a LEICA SP8 LSCM laser confocal scanning microscope, with HyD detectors HC PL APO CS2. Lasers (488nm and 561nm) were used for excitation of GFP and RFP, respectively. Metamorph 7.5 (Molecular Devices) and Image J software (National Institutes of Health, NIH, USA) were used for image analysis. 3D reconstructions were performed with Leica LAS software.

### Protoplast release assay

To evaluate the effect of chitosan on the cell wall composition of *M. oryzae*, we scored the frequency of protoplast release using cell wall degrading enzymes on chitosan grown mycelia. Guy11 was first grown in CM in the presence or absence of 0.5 and 1 mg ml^-1^ chitosan for 48h. Mycelium was then collected using Miracloth and washed twice with SDW, before being transferred to a conical tube with 40 ml of OM buffer (1.2 M MgSO_4_, 10 mM NaPO_4_; pH 5.8) in the presence of 500 mg Glucanex (Sigma) at pH 5.8. Tubes were then shaken gently to disperse hyphal clumps and then incubated for 3h at 30°C with gentle (75 rpm) shaking. Isolated protoplasts were transferred to sterile polycarbonate or polysulfonate Oakridge tubes and overlaid with an equal volume of cold ST buffer (0.6 M Sorbitol and 0.1 M Tris-HCl; pH 7.0), before centrifugation at 5,000 *g* at 4°C for 15min. Protoplasts were recovered from the OM/ST interface and transferred to a clean Oakridge tube. The tube was filled with STC (1.2 M Sorbitol, 10 mM Tris-HCl pH 7.5 and 10 mM CaCl_2_). Protoplasts were concentrated by centrifugation at 3,000 rpm/10 min/4°C, in a swinging bucket rotor, before being washed twice more with 10 ml STC. Protoplasts were resuspended in 100 µl STC buffer and counted.

### Quantification of β-1,3 glucan and chitin in cell walls of *M. oryzae*

Chitin content of mycelium was estimated by determining the amount of N-acetyl-D-glucosamine according to the method of Bowman *et al.*, (2006) with some modifications as Aranda-Martinez *et al.*, 2016. Guy11 and Δ*nox1* strains were first grown in CM for 48 h, mycelium collected, washed twice with SDW and transferred for 48h to CM in the presence or absence of 1 mg ml^-1^ chitosan. Mycelium was then collected by centrifugation, washed twice in SDW and lyophilized. Mycelium was ground in liquid nitrogen, and 30 mg per sample hydrolysed in 1 ml 6 N HCl at 110 °C for 6 h. The HCl was removed by aeration and samples resuspended in 1 ml of SDW before being centrifuged twice at 13,000 rpm for 20 min in a microfuge. A 0.5 ml aliquot of the supernatant from each sample was mixed with 0.1 ml of 0.16 M sodium tetraborate pH 9.1 and then heated at 100 °C for 3 min. After cooling, 3 ml of p-dimethylamino benzaldehyde (DMAB) solution (10% DMAB in glacial acetic acid containing 12.5% 10N HCl, diluted with 9 vol. of glacial acetic acid), was added. The mixture was then incubated for 20 min at 37°C and absorbance at 595 nm measured in a Genios(tm) Multiwell Spectrophotometer (Tecan Group AG). A standard curve was generated using 0–20 mg ml^-1^ N-acetyl-D-glucosamine (Sigma) treated as described previously.

The β-1,3-glucan content of the fungal cell wall was determined according to Shedletzky *et al*., (1997) with some modifications (Aranda-Martinez *et al.*, 2016). Mycelium of Guy11 and Δ*nox1* was transferred to CM in the presence or absence of 1 mg ml^-1^ chitosan for 48h. Mycelium were collected by centrifugation, washed twice in SDW and then hydrolysed with 0.1 M NaOH before being lyophilized. Mycelium was ground in liquid nitrogen, and 10 mg of grounded mycelium resuspended in 0.5 ml 1 M NaOH. Samples were incubated at 80°C for 30 min with 0.5 mm zirconia/silica beads (Biospec) and vortexed several times at full speed for 10 min each to favour tissues disruption. One hundred μl aliquots from each sample were then mixed with 400 μl of 1 M NaOH. Aniline blue mix (2.1 ml; 40 vol of 0.1% aniline blue, 21 vol of 1 N HCl and 59 vol of 1 M glycine/NaOH buffer; pH 9.5) was then added to each sample. These were vortexed and incubated at 50°C for 30 min then incubated for 30 min at room temperature. Fluorescence was quantified in a Jasco Model FP-6500 spectrofluorometer using 400 nm excitation and 460 nm emission wavelengths. A standard curve for β-1,3-glucan was constructed by using 0–50 mg ml^-1^ curdlan (Megazyme) dissolved in 0.1 M NaOH and also heating for 30 min at 50 °C in 1 N NaOH.

### Quantification of gene expression by qRT-PCR

Total RNA was obtained from Guy11 and Δ*nox1* cultures prepared after 48h growth in CM and further 4, 8 and 24h growth in the presence or absence of 1 mg ml^-1^ chitosan. Total RNA was isolated using Trizol reagent (Thermo Fisher Scientific) following manufacturer’s instructions. Total RNA was treated with TurboDNA free (Ambion) to remove DNA remains. First strand cDNA was then synthesized using retro-transcriptase RevertAid (Thermo Fisher Scientific) primed with oligo dT (Thermo Fisher Scientific). Gene expression was quantified using real-time reverse transcription PCR (qRT-PCR) with SYBR Green and ROX (Roche). Gene quantification was performed in a Step One Plus real-time PCR system (Applied Biosystems). Relative gene expression was estimated by the ΔΔCt methodology (Livak & Schmittgen, 2001) with three technical replicates per condition. Primers used to quantify the expression of *M. oryzae* genes in response to chitosan are shown in Table S1. Ubiquitin-conjugate enzyme (MGG_04081) and Glyceraldehyde-3-phosphate dehydrogenase (MGG_01081) genes were used as endogenous controls for all experiments (Che Omar *et al*., 2016), since their expression showed Ct stability for all conditions tested. We did not use β-tubulin and actin, commonly used as fungal housekeeping genes, because chitosan modifies their expression (Lopez-Moya *et al*., 2016). All experiments were repeated three times.

## Results

### Exposure to chitosan reduces the ability of *M. oryzae* to cause rice blast disease

In order to determine the effect of chitosan on the growth and development of the rice blast fungus *M. oryzae*, conidia were exposed to chitosan and then used to inoculate the susceptible rice cultivar CO-39. In leaf drop experiments, *M. oryzae* normally displays a large necrotic, sporulating disease lesion, as shown in Fig. 1a. However, when leaves were inoculated with *M. oryzae* in the presence of chitosan, mildly necrotic, non-sporulating lesions resulted (Fig. 1a). When conidial suspensions were used to spray-inoculate rice seedlings, exposure to chitosan (5 mg ml^-1^) significantly (p< 0.05) reduces the generation of rice blast disease lesions (Figs. 1b,c). However, spraying chitosan only in a mock inoculation does not cause any damage or response on rice leaves (Fig. 1b). Microscopic examination of rice leaf sheath preparations inoculated with *M. oryzae* conidia exposed to chitosan (5 mg ml^-1^) showed a 75% reduction in the number of cells penetrated by the fungus, compared to non-exposed, as shown in Fig. 1d. Differentiated appressoria from chitosan-treated conidia either did not penetrate epidermal rice leaf cells, or displayed a reduced invasive growth and colonisation of adjacent epidermal cells, as shown in Figs. 1e,f.

**Figure 1.**
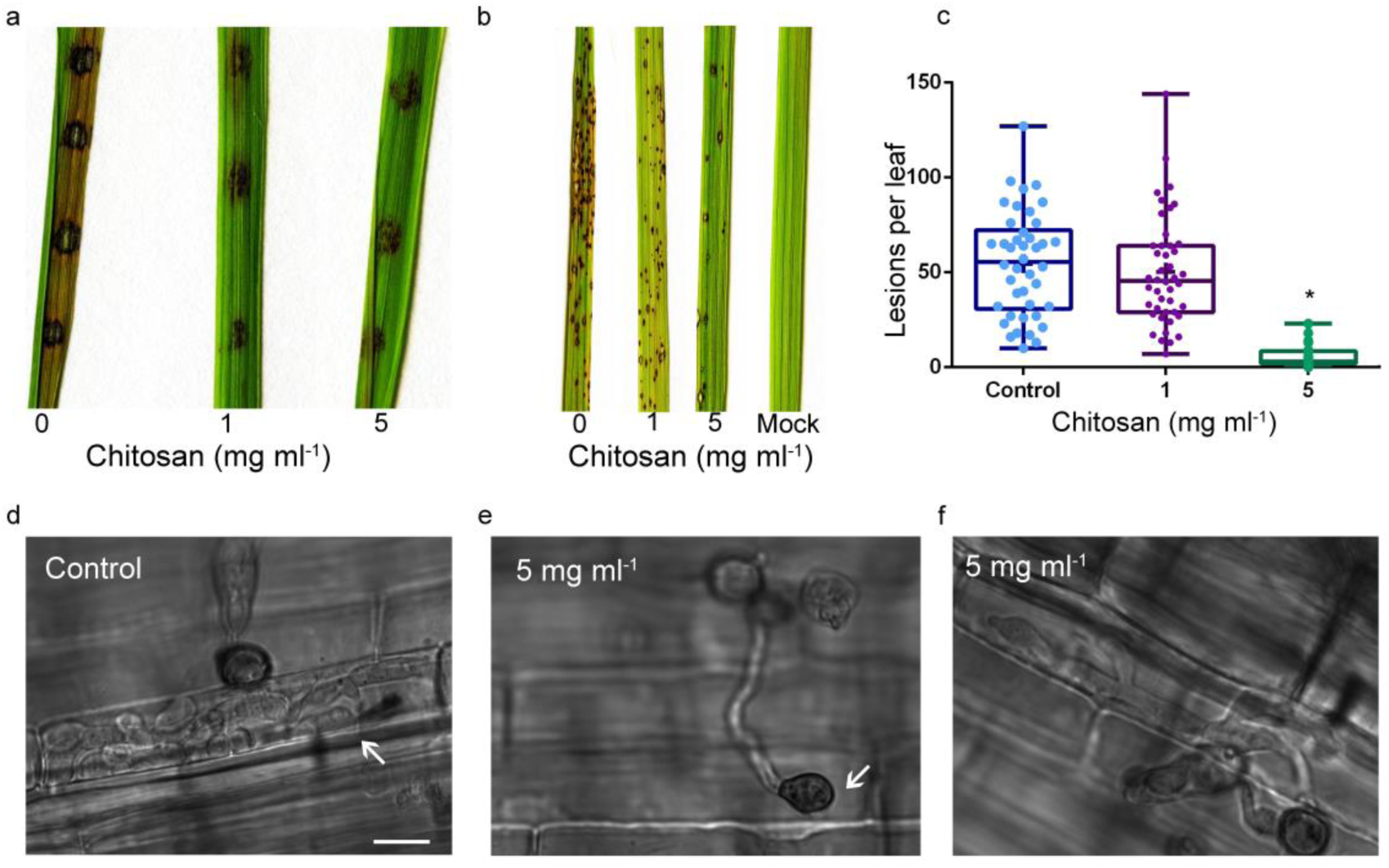
Chitosan reduces pathogenicity of *M. oryzae* on rice plants. **a)** Chitosan reduces pathogenicity of *M. oryzae* on leaf cuticle in a leaf spot bioassay after 5d. **b)** Chitosan reduces the number of lesions caused by *M. oryzae* on rice leaves in leaf spray bioassays after 5d. **c)** Chitosan significantly reduces the number of lesions per leaf after *M. oryzae* spray and incubation for 5d. Asterisk indicates significant differences. Multifactorial analysis ANOVA was used to compare treatments (p-value 0.05). **d)** Leaf sheath assay of *M. oryzae* spores colonising rice leaf cells after 30h. **e)** *M. oryzae* appressorium development under chitosan (5 mg ml^-1^) on rice leaf surface does not differentiate penetration peg and pathogenicity is inhibited after 30h. **f)** *M. oryzae* appressorium development under chitosan (5 mg ml^-1^) on rice leaf surface with reduced invasive growth compare to controls after 30h. Bar size 10µm.

### Exposure to chitosan impairs appressorium development and function in *M. oryzae*

Chitosan exposure reduces the frequency of appressorium differentiation when applied to un-germinated *M. oryzae* conidia in a concentration-dependent manner, as shown in Fig. 2. Chitosan exposure did not affect the rate of conidial germination (Fig. S1), but after 4h exposure to chitosan (0.5-2 mg ml^-1^) the number of differentiated appressoria was significantly reduced (p< 0.05). Exposure to a concentration of 5 mg ml^-1^ chitosan was sufficient to almost completely prevent appressorium differentiation (Figs. 2a,b). After 6h exposure to 5 mg ml^-1^ chitosan appressorium development was decreased by 85%. Exposure to low or intermediate concentrations of chitosan (between 0.5-2 mg ml^-1^) still cause a significant inhibition of appressorium development. Exposure to high doses of chitosan (5 mg ml^-1^) also caused an effect on appressorium shape, reducing the size of incipient appressoria (Fig. 2c and Fig. S2) and preventing formation of the appressorium melanin layer (Fig. 2d), which is essential for *M. oryzae* pathogenicity (Howard & Ferrari, 1989; Howard *et al.*, 1991).

**Figure 2.**
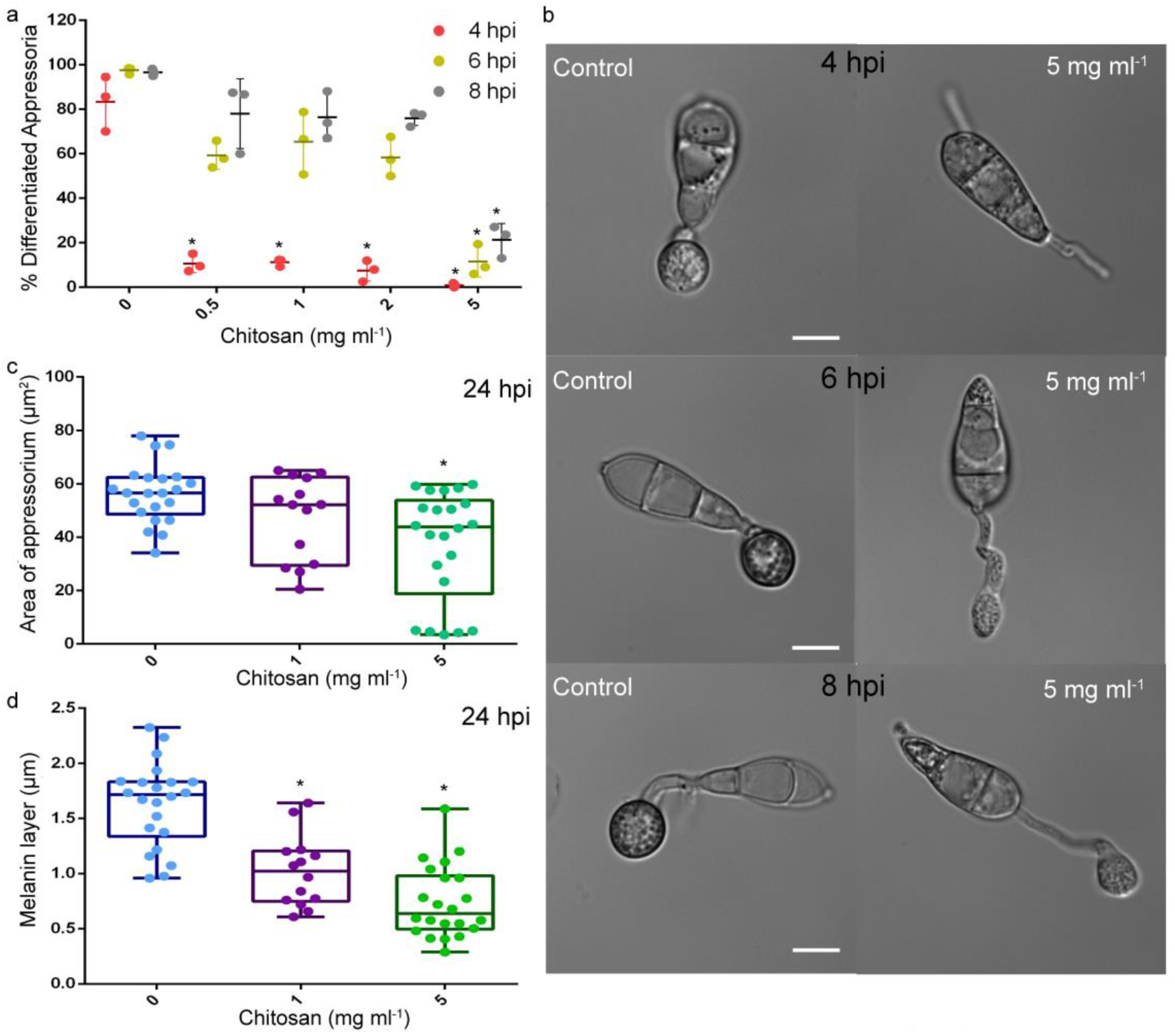
Chitosan impairs *M. oryzae* appressorium differentiation concentration-wise. **a)** Percentage of *M. oryzae* differentiated appressoria after 4, 6 and 8h in contact with (0.5-5 mg ml^-1^) chitosan. **b)** Micrographs of *M. oryzae* development after 4h, 6h and 8h with chitosan (5 mg ml^-1^) compared with untreated controls. Bar size 10 µm. **c)** Effect of chitosan on *M. oryzae* appressoria area (mean, n>20). **d)** Thickness of appressorium melanin layer differentiated under 1 and 5 mg ml^-1^ chitosan. Asterisks indicate significant differences. Multifactorial analysis ANOVA was used to compare treatments (p-value 0.05 (*)).

### Exposure to chitosan prevents septin recruitment and organisation of the appressorium pore

Appressorium re-polarisation requires recruitment and organisation of a hetero-oligomeric ring of septin GTPases at the appressorium pore (Dagdas *et al.*, 2012). In order to determine the effect of chitosan exposure on septin-dependent plant infection chitosan was applied to un-germinated conidia and the localisation of Sep4-GFP was investigated by live cell imaging, as shown in Fig. 3. *M. oryzae* appressoria normally form a large septin ring, of approximately 5.9 µm in diameter (Dagdas *et al*., 2012), as shown in Fig. 3a. Exposure to 5 mg ml^-1^ chitosan for 8h led to septin accumulation, as dense body in the centre of incipient appressoria, but no organisation into a ring structure (Fig. 3a). Similarly, when the actin-binding protein gelsolin was visualised by expression of Gelsolin-GFP, this normally formed a ring structure at the appressorium pore, which did not form following exposure of *M. oryzae* to chitosan (Fig. 3b). Consistent with the observed effect of chitosan exposure on septin organisation at the appressorium pore, the localisation of Chm1-GFP, which encodes the plasma membrane kinase which phosphorylates septins, was also affected by exposure to chitosan (Fig. 3c). Polarity determinants, such as Tea1, an F-actin–plasma membrane protein with a C-terminal actin-binding domain and N-terminal ERM domain (Gilden & Krummel, 2010) are also disorganized by exposure to chitosan (Fig. 3d). When appressoria were visualised at 24h, chitosan exposure was still sufficient to impair septin and F-actin ring organisation in *M. oryzae* appressoria (Figs. S3, S4 and S5). Quantitative analysis confirmed that chitosan treatment reduced the frequency of development of intact septin rings and the organisation of each component visualised (Fig. 3e,f,g,h).

**Figure 3.**
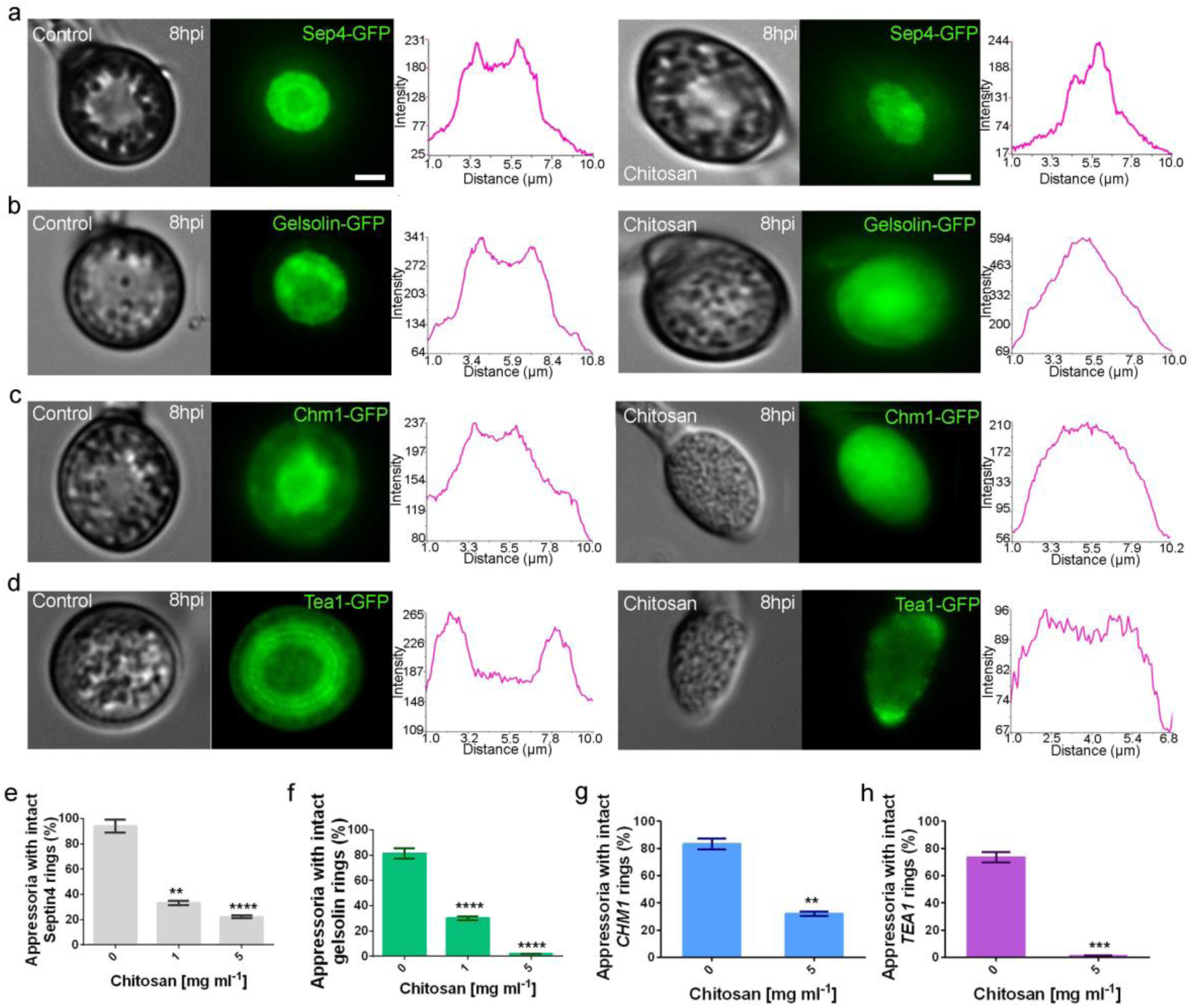
Chitosan prevents septin (SEP4) recruitment and appressorium pore organisation. **a)** Micrographs of Sep4-GFP localisation on *M. oryzae* appressorium development after 8h in the presence and absence of chitosan**. b)** Micrographs of Gelsolin-GFP localisation on *M. oryzae* appressorium development after 8h in the presence and absence of chitosan. **c)** Micrographs of Chm1-GFP localisation on *M. oryzae* appressorium development after 8h in the presence and absence of chitosan **d)** Micrographs of Tea1-GFP localisation on *M. oryzae* appressorium development after 8h in the presence and absence of chitosan. **e-h)** Linescans of (Sep4), actin (gelsolin), Chm1 and Tea1 protein fusion evaluated with no chitosan show normal ring formation after 8h. However, linescan of differentiated appressorium under chitosan did not differentiate any ring. Asterisks indicate significant differences (p-values 0.05 (*), 0.01 (**), 0.001(***) and 0.0001(****)).

In order to establish the stage at which septin organisation was perturbed by chitosan exposure, we decided to apply chitosan to *M. oryzae* germlings in a time-course experiments at 4h, 8h, 14h and 16h after conidial germination. We found that chitosan exposure at early stages of appressorium development, between 4-8h after conidial germination, prevented septin ring organisation (Fig S6; Supplementary Movie 1). There is clearly a window of activity at which chitosan exerts its effect, prior to 8h. After this point, the septin ring has already formed and chitosan exposure has no effect on its organisation (Fig. S6). We conclude that exposure to chitosan prevents septin-dependent cytoskeletal changes necessary for plant infection by the rice blast fungus.

### Chitosan permeabilises plasma membrane and perturbs the regulated synthesis of reactive oxygen species during appressorium development

The main cellular change previously reported to occur following chitosan exposure is plasma membrane permeabilization (Palma-Guerrero *et al.*, 2009; Palma-Guerrero *et al.*, 2010). To determine if this occurs upon exposure of *M. oryzae* to chitosan, the lipophilic styryl dye, FM4-64 was applied to germinating conidia in the presence or absence of chitosan (Fig. 4a). Large-scale plasma membrane permeabilization was evident based on widespread FM4-64 fluorescence detected in appressoria in the chitosan treated samples, 10min after chitosan treatment (Fig. 4b).

**Figure 4.**
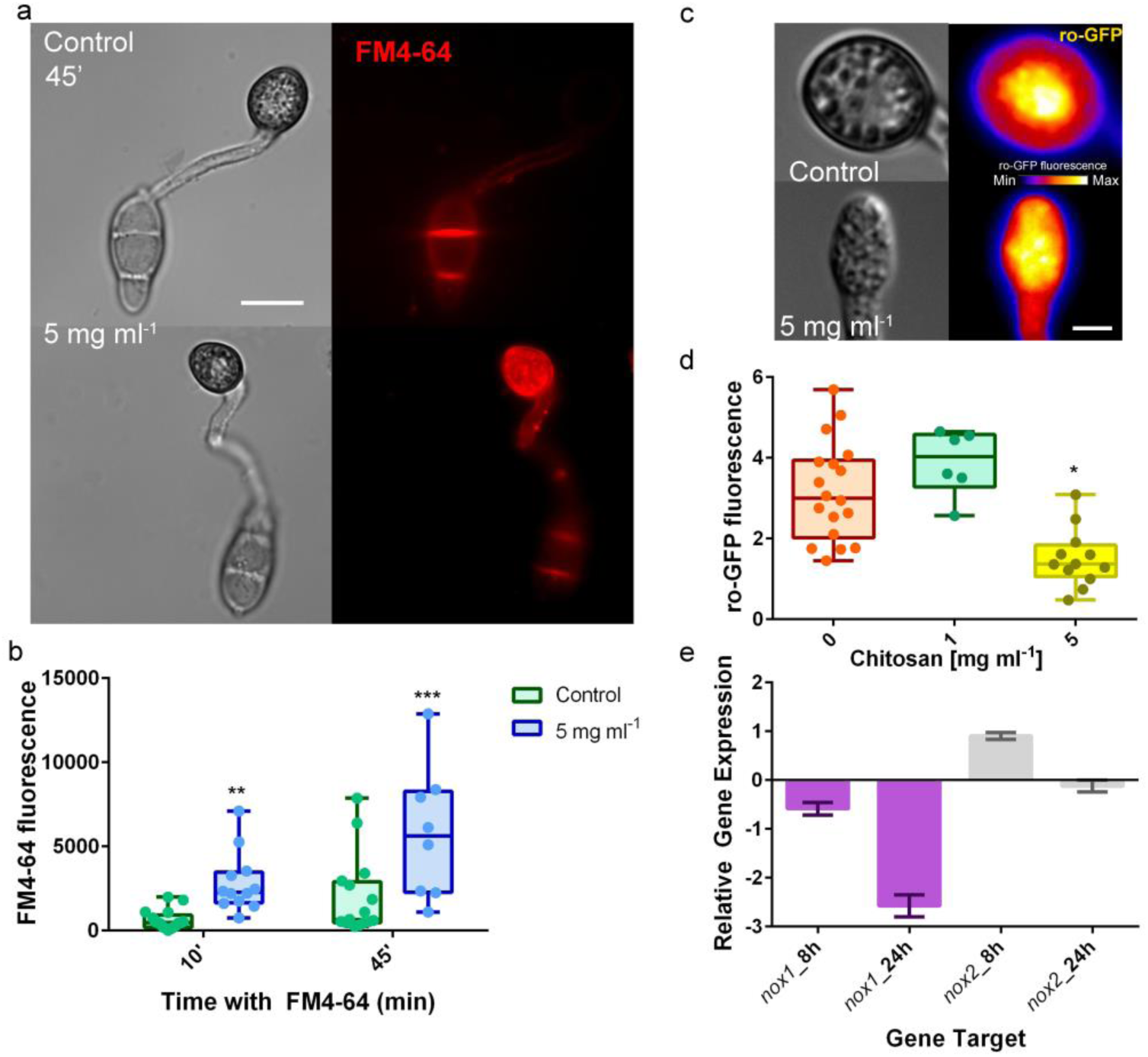
Chitosan permeabilises plasma membrane of *M. oryzae* appressorium and induces generation of reactive oxygen species. **a)** Integrity of plasma membrane on untreated 8h-old appressorium after FM4-64 application. Chitosan severely compromises plasma membrane integrity on appressoria exposed to chitosan; FM4-64 fluorescence shows high intensity within the appressorium. **b)** Average of FM4-64 fluorescence of chitosan treated appressoria and untreated controls. **c)** Fluorescence of reduced glutathione by Grx1-roGFP2 on 8h-old untreated and 5 mg ml^-1^ chitosan treated appressoria. **d)** Grx1-roGFP2 fluorescence quantitation in appressoria in the presence and absence of chitosan. **e)** NADPH oxidases (*NOX1* and *NOX2*) gene expression in Guy 11 exposed for 8h and 24h to chitosan. Chitosan early (8h) induces *NOX2* and represses *NOX1* in Guy11. However, after 24h *M. oryzae* in contact with chitosan, it drastically represses *NOX1* and slightly *NOX2.* Asterisks indicate significant differences (p-values 0.05 (*), 0.01 (**) and 0.001(***)).

The regulated synthesis of reactive oxygen species (ROS) is essential for appressorium development and necessary for septin organisation and penetration peg formation (Ryder *et al*., 2013). On the contrary, plasma membrane permeabilization by chitosan exposure leads to massive ROS generation and cell death (Lopez-Moya *et al.*, 2015). We therefore decide to use strain of *M. oryzae*, expressing Grx1-roGFP2 which detects changes in glutathione oxidation to measure ROS accumulation in *M. oryzae* appressoria. We observed a significant (p< 0.05) reduction in Grx1-roGFP2 fluorescence in appressoria following exposure to 5 mg ml^-1^ chitosan for 8h (Fig. 4c and 4d). We also observed in Guy11 elevated expression of the *NOX2* NADPH oxidase, but a small decrease in *NOX1* expression in response to chitosan treatment (Fig. 4e). To investigate this response further, we then examined the expression of *NOX2* in a Δ*nox1* mutant. We observed increased *NOX2* expression in the Δ*nox1* mutant in the presence of chitosan (Fig. S7). We conclude that chitosan exposure increases ROS generation and permeabilizes plasma membrane of the rice blast fungus.

### Sensitivity to chitosan requires Nox1-dependent NADPH oxidase activity

Given the effect of chitosan exposure to ROS generation and membrane permeability in *M. oryzae*, we decided to investigate the phenotype of chitosan treatment on Δ*nox1*, Δ*nox2* mutants, lacking the respective catalytic sub-units of the NADPH oxidases (Egan *et al*., 2007). To carry out this experiment, we exposed each mutant to 0.5-1 mg ml^-1^ chitosan and measured their vegetative growth in liquid CM cultures, based on dry weight. Then, we observed that chitosan exposure-initiated accumulation of melanin in culture filtrates of the wild-type strain Guy 11 and in the Δ*nox2* mutant, albeit only in response to higher chitosan concentrations (Fig. 5a). Strikingly, however, Δ*nox1* mutants did not secrete excess melanin (Fig. 5a). Consistent with the difference in appearance of liquid cultures, we found that Δ*nox1* mutants were more resistant to chitosan exposure, than Guy11 and Δ*nox2* mutants, which both showed significant reductions in dry weight following exposure to chitosan compared to the Δ*nox1* mutant (Fig. 5b). We observed that chitosan treatment of a Δ*nox1nox2* double mutant did not cause any reduction in biomass and cultures continued to grow very well (Fig. 5b). This suggests that absence of the Nox1 catalytic sub-unit of the NADPH oxidase complex leads to enhanced tolerance to chitosan treatment and that this remains unaffected by further loss of Nox2.

**Figure 5.**
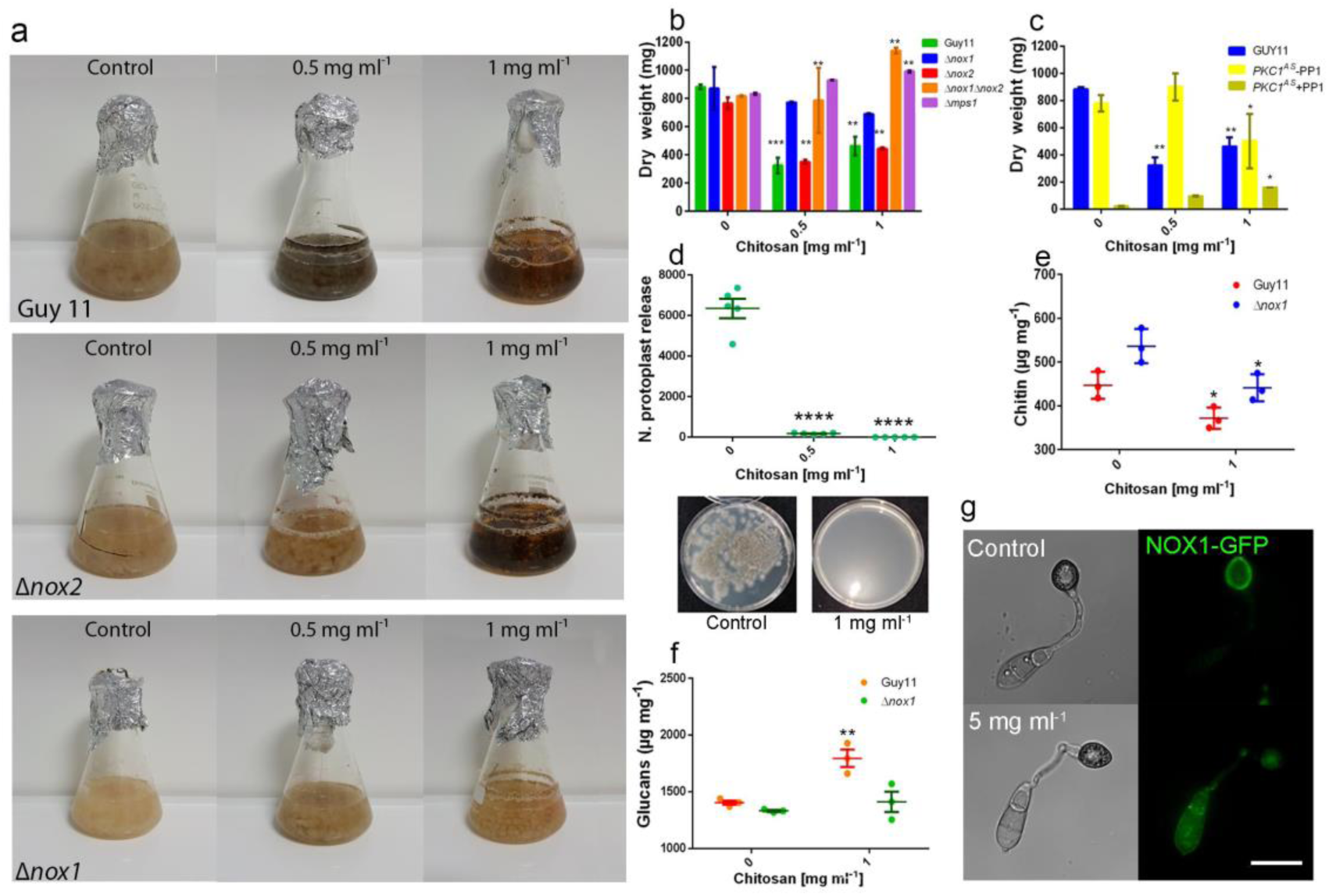
*NOX1* and CWI pathway are essential for *M. oryzae* sensitivity to chitosan. **a)** Mycelium from Guy11, Δ*nox1*, Δ*nox2*, and Δ*nox1nox2* mutants was inoculated onto complete medium containing chitosan at a concentration 0.5 and 1 mg ml^-1^ and incubated at 24°C for 2 d with a 12h light/dark cycle. **b)** Δ*nox1*, but specially Δ*nox1nox2*, show high tolerance for growing with chitosan in CM medium indicating the key role of Δ*nox1* in sensitivity of *M. oryzae* to chitosan. Δ*mps1* is also able to grow under high doses of chitosan indicating. **c)** Guy 11 and *Pkc*^*AS*^ in absence of NA-PP1 showed a significant decrease of biomass growing with chitosan respect to untreated control. However, when *Pkc*^*AS*^ is growing with chitosan and NA-PP1 is added in the media, this strain showed increased tolerance to chitosan. **d)** Guy 11 mycelium grown in the presence of chitosan was more resistant to cell wall degrading enzymes, showing significantly less protoplast release. **e)** Guy11 show less amount of chitin in its cell wall than Δ*nox1* indicating the key role of *NOX1* in the cell wall biosynthesis. Chitosan propitiates in both Guy11and Δ*nox1* a significantly reduced amount of chitin in their cell walls. **f)** Guy11and Δ*nox1* shows similar amount of glucan in their cell walls. However, exposure of these strains to chitosan increases glucan content in Guy 11, and to a less extent in Δ*nox1* **g)** Micrograph of Nox1-GFP localisation on *M. oryzae* appressorium development after 8h en the presence and absence of chitosan. Multifactorial analysis ANOVA was used to compare treatments (p-values 0.05 (*), 0.01 (**), 0.001(***) and 0.0001(****)).

### Sensitivity to chitosan requires the protein kinase C-dependent cell wall integrity pathway in *M. oryzae*

The Δ*nox1* mutant has previously been shown to be more resistant to calcofluor white (Egan *et al.*, 2007), suggesting that Nox1 is critical for correct cell wall biosynthesis. This is consistent with its role in penetration peg elongation and invasive fungal growth (Ryder *et al.*, 2013). We reasoned that the resistance to chitosan treatment shown by Δ*nox1* mutants may suggest that the uptake and/or fungicidal role of chitosan may be dependent on correct organisation of the fungal cell wall. To test this idea, we evaluate the effect of chitosan on Δ*mps1* mutants, impaired in function of the cell wall integrity pathway (Xu *et al.*, 1998). The Mps1 MAP kinase is important for hyphal growth, conidiation and appressorium function and mutants show defects in their regulation of cell wall biogenesis (Xu *et al*., 1998). We observed that Δ*mps1* mutants showed enhanced resistance to chitosan exposure, growing as well as a Δ*nox1nox2* double mutant in the presence of high concentrations of chitosan (Fig. 5b).

The cell wall integrity pathway is regulated by protein kinase C (Levin 2011) and in *M. oryzae, PKC1* is an essential gene required for cell viability (Penn *et al.*, 2015). To test whether chitosan sensitivity requires Pkc1-dependent signalling, we therefore used an analogue-sensitive mutant of Pkc1, which is sensitive to the ATP analogue 4-amino-1-tert-butyl-3-(1’-naphthyl) pyrazolo[3,4-d] pyrimidine (NA-PP1). This was generated by mutation of the gatekeeper residue of the ATP-binding pocket of Pkc1 and targeted allelic replacement to provide a mutant in which Pkc1 activity can be specifically inhibited by application of 1NA-PP1 (described by Penn *et al*., 2015). This mutant normally fails to grow in the presence of 1NA-PP1. We incubated the *pkc1*^*AS*^ mutant in the presence or absence of chitosan and 1 µM NA-PP1. The lethal effect of 1NA-PP1 was partially remediated by the presence of chitosan, allowing some fungal growth to occur, as shown in Fig. 5c. This result is consistent with absence of the Pkc1-dependent protein kinase C pathway leading to enhanced resistance to chitosan. However, the partial remediation of the lethal effect of losing Pkc1 activity by chitosan may also point to a more direct role for chitosan in stabilising the cell wall when it is affected in composition by impairment of the cell wall integrity signalling pathway.

To investigate the effect of chitosan on cell wall composition in *M. oryzae*, we carried out a protoplast release assay, which evaluates the sensitivity of the cell wall to cell wall degrading enzymes. We treated fungal mycelium grown in the presence or absence of chitosan (0.5 and 1 mg ml^-1^), with Glucanex (Sigma Aldrich), a commercial preparation of glucanases and chitinases, to release protoplasts. We observed that mycelium grown in the presence of chitosan was more resistant to cell wall degrading enzymes, showing significantly (p< 0.0001) less protoplast release (Fig. 5d and Fig. S8).

We then determined the chitin and glucan content of cells walls of mycelium of *M. oryzae*, grown in the presence or absence of chitosan (Fig. 5e,f). We observed that chitosan exposure led to a significant decrease in chitin content and increase in glucan content of fungal cell walls of the wild-type strain Guy11. Interestingly, when we repeated this assay in a Δ*nox1* mutant, we observed an elevated chitin content and reduced glucan content in the mutant, when grown normally in the absence of chitosan. Chitosan treatment led to a decrease in chitin content of a Δ*nox1* mutant, but these levels were still the same as in those in Guy11 grown normally. Furthermore, there was little overall effect on glucan content in the Δ*nox1* mutant when exposed to chitosan (Fig. 5e,f). These observations suggest that the Δ*nox1* mutant may be resistant to chitosan due to its enhanced glucan/chitin cell wall ratio. Consistent with the importance of Nox1 function in the response to chitosan, live cell imaging experiments of a *M. oryzae* strain expressing Nox1-GFP revealed misslocalisation of the NADPH oxidase after exposure to chitosan. Nox1-GFP is normally located at the appressorium cortex in 24h appressoria. However, in appressoria developed in the presence of a high chitosan dose (5 mg ml^-1^), Nox1-GFP was located in the central appressorium vacuole (Fig. 5g). When considered together, our observations suggest that the cell wall integrity pathway is critical to the fungicidal activity of chitosan and this may be linked to the glucan/chitin ratio of fungal cell walls.

### Exposure to chitosan causes repression of cell wall integrity pathway gene expression

To characterise the link between the cell wall integrity pathway and the response of *M. oryzae* to chitosan, we studied expression of genes encoding components of the cell wall integrity pathway in mycelium grown in the presence or absence of chitosan. We observed that chitosan exposure led to repression in the expression of *MPS1, PKC1, RHO1* and *SWI6* in Guy 11 (Fig. 6). In all cases the effect was most pronounced after 8h exposure, but some repression was maintained even after 24h exposure to chitosan. When the same genes were analysed in a Δ*nox1* mutant in the presence or absence of chitosan, we observed that chitosan exposure led to elevated expression of *PKC1, MPS1, RHO1* and *SWI6* after 24h exposure (Fig. 6). *MPS1* expression was initially reduced in the Δ*nox1* mutant when exposed to chitosan for 8h, but then showed elevated expression by 24h. The overall pattern of gene expression suggests that chitosan treatment normally leads to repression of the cell wall integrity pathway. By contrast, the absence of the Nox1 NADPH oxidase prevents this repression mediated by chitosan exposure and instead leads to elevated expression. Taken together, these results are consistent with cell wall integrity pathway function being essential for the fungicidal activity of chitosan.

**Figure 6.**
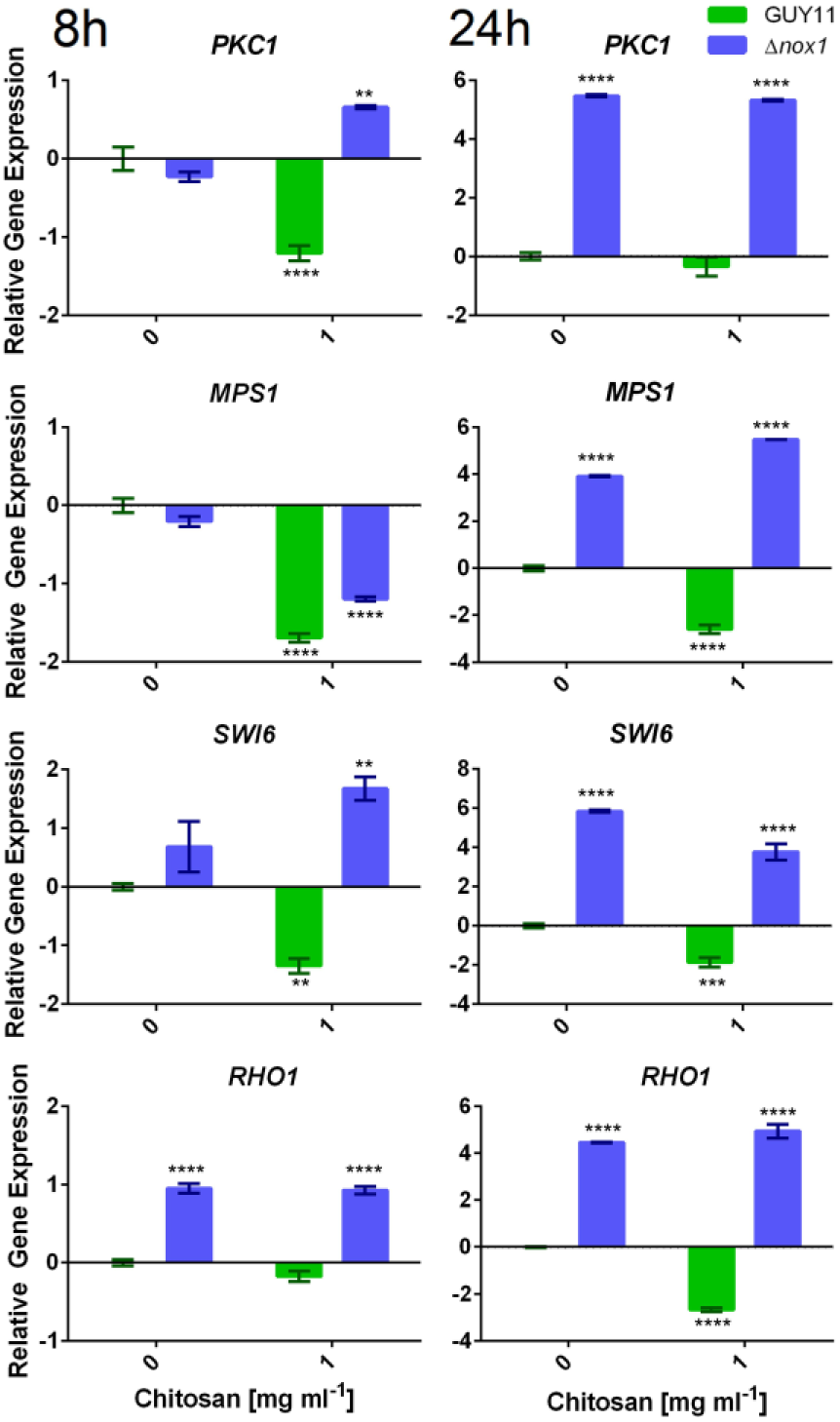
Chitosan represses *PKC1* pathway through *NOX1* activity. Exposure of Guy11 to chitosan for 8h and 24h propitiates *PKC1, MPS1, SWI6* and *RHO1* repression through NOX1 activity. Δ*nox1* show overexpression of the main CWI pathway genes (*PKC1, MPS1, SWI6* and *RHO1*) after 8h. This would indicate that NOX1 acts by repressing *M. oryzae* CWI pathway gene expression in response to chitosan. Multifactorial analysis ANOVA was used to compare treatments (p-values 0.05 (*), 0.01 (**), 0.001(***) and 0.0001(****)).

## Discussion

Chitosan has considerable potential as a naturally occurring anti-fungal agent. It can be readily produced by partial deacetylation of chitin, one of the most common biopolymers, a constituent of the cell walls of fungi, and the exoskeletons of arthropods and crustaceans (Allan & Hadwiger, 1979). Commercially, chitosan can be produced in large quantities, for example, from crab shell waste (Kumar, 2000). In addition to being a highly toxic cationic inhibitor of the growth of many fungal species, including important plant pathogens, chitosan is not toxic to mammals, including humans (Lopez-Moya *et al.*, 2015), or to plants leaves, where it can act instead as a plant-defence inducing compound (Trotel-Aziz *et al.*, 2006). Chitosan therefore has considerable potential to treat plant diseases, but its mode-of-action is still not completely clear.

In this study we set out to determine the effects of chitosan treatment on the rice blast fungus *M. oryzae*, one of the world’s most devastating crop pathogens, which causes very substantial losses to the annual rice harvest (Wilson & Talbot, 2009; Martin-Urdiroz *et al.*, 2016). The infection mechanism of *M. oryzae* has also been intensively studied making it an excellent model for understanding how chitosan might act in the context of perturbing plant infection by a phytopathogenic fungus.

First of all, we have demonstrated that chitosan is able to inhibit the growth of *M. oryzae* and its ability to cause rice blast disease. It is known that chitosan forms part of the differentiated cell wall of *M. oryzae* during appressorium-mediated plant infection (Geoghegan & Gurr, 2016), but the fungicidal action of exogenously applied chitosan to *M. oryzae* has not previously been shown. We observed that chitosan blocks appressorium-mediated plant infection by the fungus and is able to prevent organisation of the septin ring, that is necessary for F-actin re-organisation at the appressorium pore. Four core septins, Sep3, Sep4, Sep5 and Sep6 are essential for generating a hetero-oligomeric ring structure that bounds the appressorium pore and is essential for plant infection (Dagdas *et al.*, 2012). Formation of this ring structure is pivotal to penetration peg development and therefore rupture of the rice cuticle. The septin ring acts as a scaffold to organise F-actin at the point of infection and as a lateral diffusion barrier to hold polarity determinants, such as Tea1, Cdc42, and Las17 at the centre of the pore, from where formation of the penetration peg occurs (Dagdas *et al*., 2012; Ryder *et al*., 2013). Chitosan prevents these developmental changes from occurring and polarity of the appressorium is thus impeded, preventing plant infection. Septins are well known regulators of polarity and fungal morphogenesis in filamentous fungi, such as *Aspergillus nidulans* and *Neurospora crassa* (Berepiki & Read, 2013) and yeasts such as *S. cerevisiae* and *Schizosaccharomyces pombe* (Gladfelter *et al.*, 2005; Hernandez-Rodriguez & Momany, 2012). Consistent with this effect on septin recruitment and organisation, previous transcriptional profiling experiments in *N. crassa* show that chitosan exposure represses expression of the core septin-encoding genes *CDC10, CDC11*, and *CDC12* (Lopez-Moya *et al.*, 2016). The inhibition of septin recruitment was also shown to impair F-actin organisation during appressorium differentiation and the localisation of Tea1 and Chm1. This is consistent with chitosan preventing appressorium function due to an inability to carry out the rapid septin-dependent, actin polymerisation at the base of the infection cell, necessary for re-polarisation. In *N. crassa*, proteins associated with actin polymerisation proteins are also repressed by chitosan (Lopez-Moya *et al.*, 2016). These results support the hypothesis that one of the consequences of chitosan exposure is to disrupt cytoskeletal organisation.

To investigate the primary mode-of-action of chitosan we decided to evaluate its effect on the plasma membrane and cell wall. Previous studies have shown that chitosan exposure causes plasma membrane permeabilisation (Palma-Guerrero, *et al.*, 2009; Jaime *et al.*, 2012, Lopez-Moya, *et al.*, 2015). It is known that chitosan permeabilises plasma membrane in an ATP-dependent manner in *N. crassa* (Palma-Guerrero, *et al.*, 2009) and that its toxicity is dependent on membrane fluidity. A fatty acid desaturase mutant, *Δods*, showing reduced plasma membrane fluidity exhibited increased resistance to chitosan (Palma-Guerrero *et al.*, 2010). Chitosan resistant fungi, such as the nematophagous fungus *Pochonia chlamydosporia*, also showed reduced polyunsaturated fatty acid content in the plasma membranes (Palma-Guerrero *et al.*, 2010). We observed that chitosan exposure led to plasma membrane permeabilization in *M. oryzae*, consistent with these previous studies. Permeabilization and disruption of membrane integrity may lead to the changes in septin organisation observed, given the importance of the plasma membrane cytoskeleton scaffolding (Porter & Day, 2015).

It is known that membrane-localised NADPH oxidases are key to regulating the recruitment and organisation of septins during plant infection by *M. oryzae* (Egan *et al.*, 2007 and Ryder *et al*., 2013). We observed that chitosan exposure causes massive ROS synthesis, consistent with its effects on septin recruitment. In this sense, we also observed elevated expression of *NOX2* by chitosan treatment. We also found that a Δ*nox1* mutant, which lacks one of the catalytic sub-units of NADPH oxidase, is more resistant to chitosan treatment. Previously it has been reported that Nox1 is required for cell wall organisation in *M. oryzae* as Δ*nox1* mutants show greater resistance to calcofluor white (Egan *et al.*, 2007). It is also known that Nox1 is necessary for penetration peg elongation and is therefore likely to be a key regulator of cell wall biosynthesis. We observed that Δ*nox1* mutants showed a higher chitin content than an isogenic wild-type strain of *M. oryzae* and this may have contributed to its resistance to chitosan, as exposure to chitosan appears to deplete the fungus of chitin in its cell walls. The imbalance in chitin and glucan content caused by chitosan exposure may be an important element of its toxicity and may also be associated with the ability of chitosan to traverse the wall effectively in order to bind to its primary target, the plasma membrane (Palma-Guerrero *et al*., 2009; Palma-Guerrero *et al*., 2010; Lopez-Moya *et al.*, 2015; Lopez-Moya *et al.*, 2016; Aranda-Martinez *et al.*, 2016).

The importance of the fungal cell wall in conditioning the response of *M. oryzae* to chitosan exposure led us to examine mutants in key components of the cell wall integrity pathway. The cell wall integrity pathway is well known to mediate the response to cell wall stress and is therefore associated with osmotic stress adaptation and xenobiotic stresses, including drug treatments (Bahn *et al.*, 2007; LaFayette *et al.*, 2010; Penn *et al.*, 2015). We found that chitosan exerts its toxicity towards *M. oryzae* through a process dependent on a functional cell wall integrity pathway. A Δ*mps1* mutant lacking the cell wall integrity MAP kinase, was more tolerant to chitosan than the wild-type strain. *MPS1, PKC1, SWI6* and *RHO1* genes were all repressed in response to chitosan treatment. *RHO1* controls cell wall synthesis and aggregation of actin cables in *M. oryzae* (Fu *et al.*, 2018) and its down-regulation in response to chitosan is consistent with the defects in F-actin organisation observed after drug treatment. Protein kinase C, which acts as a major control point for operation of the cell wall integrity pathway is essential in *M. oryzae*, but we found that a conditionally lethal, analogue-sensitive mutant of *PKC1* (Penn *et al.*, 2015), could be partially remediated for growth in the presence of chitosan. This result strongly suggests that chitosan can partially serve to stabilise cell wall integrity in the absence of the PKC signalling pathway in a way that allows some fungal growth to occur. It is clear, however, that when the PKC cell wall integrity pathway is fully operational, chitosan can permeabilise the cell membrane, affect NADPH oxidase functions, disorganize septin and disrupting the actin cytoskeleton. The role of the Nox1 NADPH oxidase in regulating cell wall biogenesis is also clear, because its absence renders the fungus much less sensitive to chitosan, up-regulates genes encoding components of the PKC cell wall integrity pathway in the presence of chitosan, and leads to an elevated chitin cell wall content. This change in cell wall composition explains the previously reported resistance of Δ*nox1* mutants to calcofluor white (Egan *et al.*, 2007) and reveals the importance of cell wall function to the ability of chitosan to inhibit fungal growth.

Chitosan is therefore a potential means by which rice blast disease could be controlled in future, given its ability to prevent leaf infection at a very early stage, prior to cuticle penetration. Furthermore, because chitosan is likely to act at multiple sites in conditioning membrane permeabilization, the chances of selecting specific resistant mutants is likely to be very low. Results here also show how increased tolerance to chitosan, which might emerge by perturbation of cell wall function is unlikely to result in mutants that could survive under field conditions because of the essential nature of cell wall composition and its regulation to growth and development. In conclusion, we show new data on the mode of action of chitosan useful to control a plant pathogen threatening global food security. We propose chitosan as a useful tool to modify fungal morphogenesis, gene expression and pathogenicity. Our results are, therefore, of paramount importance for developing chitosan as a natural antifungal to control rice blast.

## Acknowledgements

This work was supported by AGL 2015 66833-R grant from the Spanish Ministry of Economy and Competitiveness and European H2020 Project MUSA-727624. We would like to thank Dr Nuria Escudero (Microomics Systems S.L.) and Msc Neftaly Cruz-Meireles (TSL) for their technical support. Authors would like to thank Dr Mick Kershaw (University of Exeter) for his contribution for manuscript editing.

## Author Contribution

F. L-M: design and performance of the research, data collection and data analysis, writing the manuscript., M. M-U.: performance of the research and technical support and data interpretation, M. O-R.: performance of the research and technical support and data interpretation, M. F.: ro-GFP results analysis and other data interpretation, G.L.: performance of the research and technical support with confocal imaging and data interpretation, L.V. L-Ll.: design of the research, performance of the research, data interpretation, writing the manuscript., N.J.T.: design of the research, data interpretation, writing the manuscript.

