## Supplementary Figures for "Chitosan inhibits septin-mediated plant infection by the rice blast fungus *Magnaporthe oryzae* in a Protein Kinase C and Nox1 NADPH oxidase-dependent manner"

**Figure S1.** ***M. oryzae*** **germination with chitosan.** Chitosan did not affect *M. oryzae* germination after 2h exposure (n>50). Bars indicate standard deviation. This experiment was replicated 3 times.

**Figure S2. Morphometrical analysis of the effect of chitosan on appressorium development. a)** Micrograph of 24h-old appressoria developed without chitosan. Lines indicate the morphometric parameters measured. Yellow line (D1) shows distance (µm) between the diameter and the base of the appressorium. Red line (D2) shows distance (µm) diameter perpendicular to D1. Green line (ML) shows the thickness (µm) of the melanin layer. Blue line defines the appressorium area. **b)** Micrograph of 24h-old *M. oryzae* appressorium developed under 1mg $ml^{-1}$ chitosan. **c)** Micrograph of 24h-old appressorium developed under 5mg ml-1 chitosan. **d)** Average of both D1 and D2 appressorium diameters. Asterisks indicate significant differences ($p< 0.05$). Bar size 10µm.

**Figure S3. Chitosan inhibit septin (Sep4) organisation by inhibiting septin ring formation in *M. oryzae*. a)** Micrograph of *M. oryzae* appressorium development after 4h, 6h and 24h with no chitosan**. b)** *M. oryzae* appressorium development after 4h, 6h and 24h under 5 mg $ml^{-1}$ chitosan. **c)** Linescan of differentiated appressorium with no chitosan show septin ring formation after 24h. **d)** Linescan of differentiated appressorium under chitosan does not show septin rings. Bar size 10µm.

**Figure S4. Chitosan blocks gelsolin organisation by inhibiting pathogenic actin ring formation in *M. oryze*. a)** Micrograph of *M. oryzae* appressorium development and gelsolin organisation after 4h, 6h and 24h with no chitosan**. b)** *M. oryzae* appressorium development and gelsolin after 4h, 6h and 24h under 5 mg $ml^{-1}$ chitosan. **c)** Linescan of differentiated appressorium with no chitosan show normal actin associated protein gelsolin ring formation after 24h. **d)** Linescan of differentiated appressorium under chitosan do not show actin associated protein gelsolin rings. Bar size 10µm.

**Figure S5. Chitosan affects organisation and localisation of Chm1-GFP and Tea1-GFP in *M. oryzae*.** Chitosan impairs Chm1 localisation on a pore in the centre of the appressorium after 24h. Application of chitosan drastically mislocalises Chm1 and a dense body is generated. Linescan shows Chm1 misslocalisation around all the appressorium. Untreated appressorium organises Chm1 into a pore. Chitosan also impairs Tea1 localisation, drastically misslocalises Tea1 and a dense body is generated after 24h. Linescan also shows Tea1 disorganization after 24h. Bar size 10µm.

**Figure S6. Chitosan applied at early stages of appressorium development seriously impairs septin organisation in *M. oryzae*.** Untreated appressorium organises septin in a ring after 24h. Application of **c**hitosan applied on 4h-old appressoria drastically misslocalise the septin a dense body of the appressorium after 24h. In addition, application of chitosan on 8h-old appressorium also inhibits septin ring organisation. Surprisingly, application of Chitosan on 14h-old appressorium partially affects septin ring organisation after 24h. Similar results were found when chitosan was applied on 16h-old appressoria. Number of intact rings are drastically reduced when chitosan is applied at early stages of appressorium differentiation (n>15). Asterisk indicates significant differences ($p < 0.05$). Bar size 10µm.

**Figure S7. Chitosan increases *NOX2* expression in the *M. oryzae* Δ*nox1* mutant.** Δ*nox1* propitiates an over expression of *NOX2* indicating a compensatory effect between NADPH oxidases. When chitosan is applied *NOX2* is still more induced in Δ*nox1* probably generating ROS inside of the fungal cells. These results are agreed with the key role of Nox1 in *M. oryzae* response to chitosan.

**Figure S8. *M. oryzae* mycelium grown in the presence of chitosan is resistant to cell wall degrading enzymes**. Mycelium grown in the presence of chitosan was more resistant to cell wall degrading enzymes, showing significantly ($p < 0.0001$) less protoplast release. It is observed because in the chitosan-treated mycelia is not formed the typical layer o protoplasts which is visible in the untreated controls

**Movie S1. Chitosan applied at early stages (8h) of *M. oryzae* appressorium development blocks septin organisation**

**Table S1. Primer Sequences**

| Primer Sequence (5'→3') | Primer Name | Size (bp) |
|---|---|---|
| ACGGTGTCGAGTTCCTCAAG | NOX1_F | 85 |
| ACCGAGTTGAGCACGATGTT | NOX1_R1 | |
| ATCCCTCTCAGCGTGTACCT | BKC1_F1 | 96 |
| GCTGAGAGCCAGACATTGGT | BKC1_R1 | |
| ACCCACGACTTCCTCTACGA | SEP5_F1 | 62 |
| CCACCATCAACACTCCTGCT | SEP5_R1 | |
| TTTGCGACTTTGGTCTTGCG | MPS1_F1 | 89 |
| GTACCACCTGGTAGCCACAT | MPS1_R1 | |
| TATGAGCCGTGGTGCTTCTG | NOXR_F1 | 98 |
| TGGTACGTTCGTAGCGGTCATT | NOXR_R | |
| ACGCTGCAGATCAAGGTCAA | CHM1_F1 | 88 |
| TCACTCCTCCCATACCAGGG | CHM1_R1 | |
| TATCGCCCGTGAGATTCGTG | NOX2_F1 | 64 |
| GCTGGGGTGCTGGATAACTT | NOX2_R1 | |
| ATTGTTGACACGCCAGGCTA | SEP4_F1 | 94 |
| GGTAGGCAGAGTGCTGATCC | SEP4_R1 | |
| ACGTCGAGTTGGCACTATGG | RHO1_F1 | 80 |
| GTGAGAGTCGGGGTAGGACA | RHO1_R1 | |
| AGCAAGGGTATGGTCAACCG | PKC1_F1 | 99 |
| GCTCGAGGTTAAACCAGCCT | PKC1_R1 | |
| ATGAACGATGGCATGTCGGA | SWI6_F1 | 70 |
| TGTCGAGGTCTCTCGGATGT | SWI6_R1 | |
| CTGCCATCTTCCGTGGAAAGG | TUB_F | 86 |
| GACGAAGTACGACGAGTTCTTG | TUB_R | |
| CATCTTAACGTCGTCGTCATC | EF1_F | 62 |
| AGTGGCCGGTAGTCGTGG | EF1_R | |
| ATCCTAATGTCTACCCGAG | Ubiq_F | 85 |
| GATGCGTGTTCGTAGTGG | Ubiq_R | |
| CAAGTACGCCAAATACATGC | GAPDH_F | 96 |
| TTGCCGTTGACGACCAGG | GAPDH_R | |

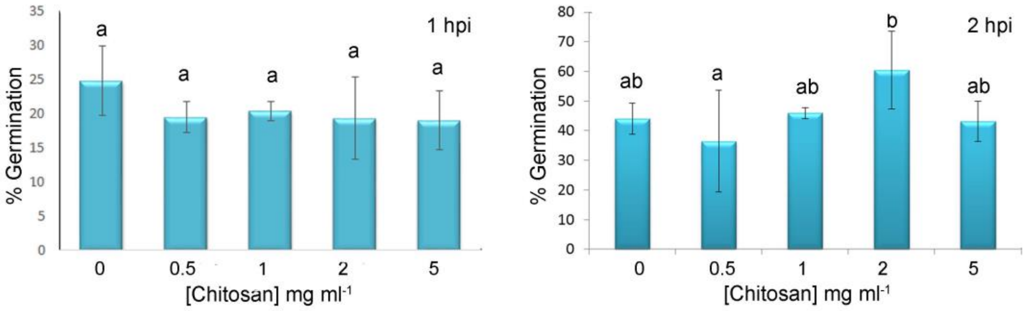

1 hpi
% Germination
0
5
10
15
20
25
30
35
a
a
a
a
a
0
0.5
1
2
5
[Chitosan] mg ml-1
2 hpi
% Germination
0
10
20
30
40
50
60
70
80
ab
a
ab
b
ab
0
0.5
1
2
5
[Chitosan] mg ml-1


Figure S1

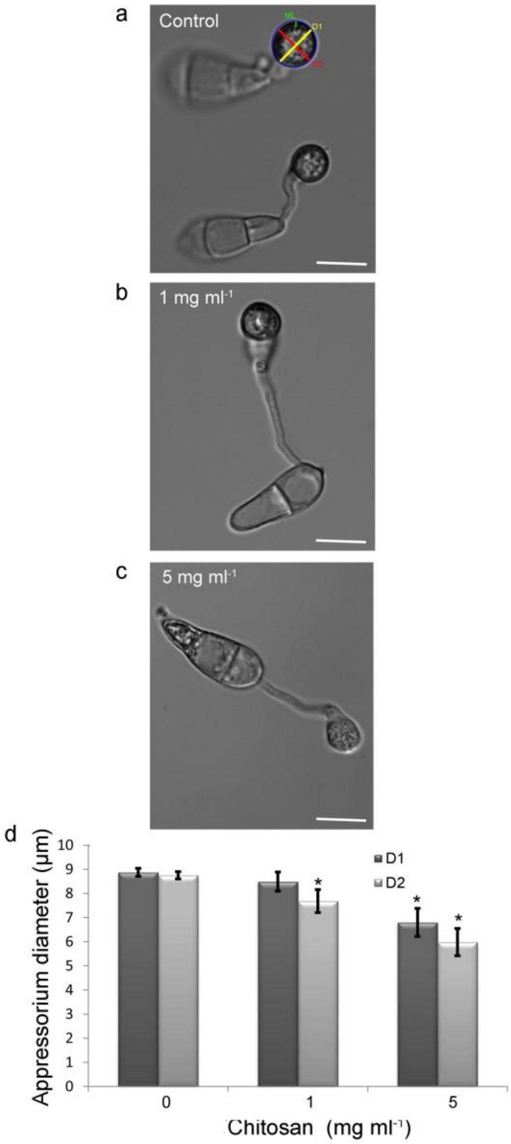
a
Control
b
1 mg ml-1
c
5 mg ml-1
d
Appressorium diameter (μm)
10
9
8
7
6
5
4
3
2
1
0
D1
D2
*
0
1
5
Chitosan (mg ml-1)

Figure S2

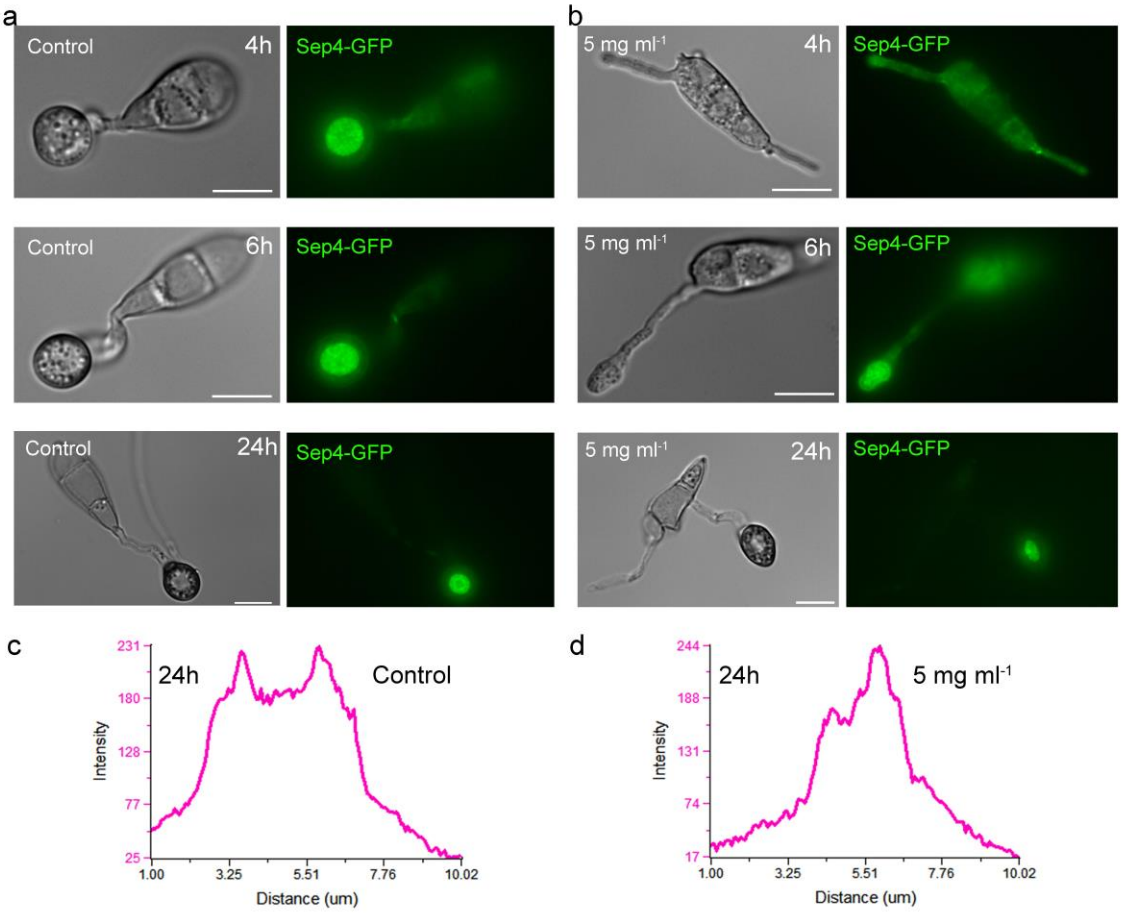
a
Control
4h
Sep4-GFP
Control
6h
Sep4-GFP
Control
24h
Sep4-GFP
b
5 mg ml-1
4h
Sep4-GFP
5 mg ml-1
6h
Sep4-GFP
5 mg ml-1
24h
Sep4-GFP
c
24h
Control
Intensity
231
180
128
77
25
1.00
3.25
5.51
7.76
10.02
Distance (um)
d
24h
5 mg ml-1
Intensity
244
188
131
74
17
1.00
3.25
5.51
7.76
10.02
Distance (um)

Figure S3

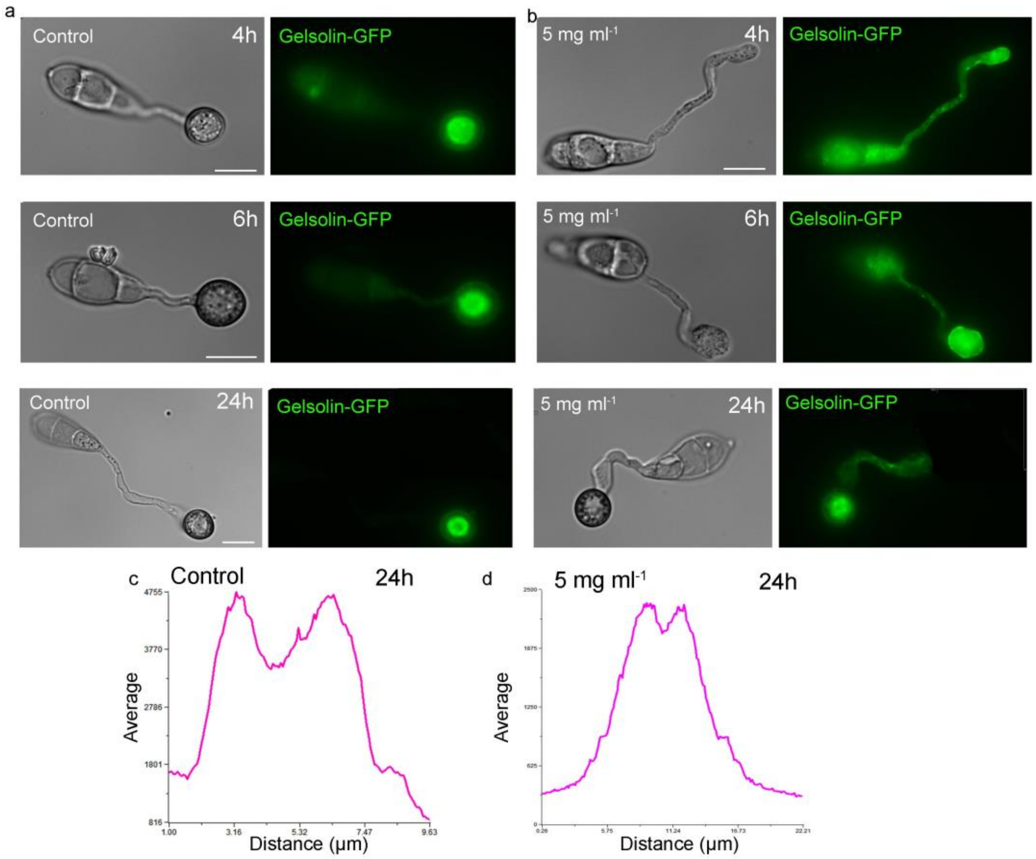
a
Control
4h
Gelsolin-GFP
Control
6h
Gelsolin-GFP
Control
24h
Gelsolin-GFP
b
5 mg ml-1
4h
Gelsolin-GFP
5 mg ml-1
6h
Gelsolin-GFP
5 mg ml-1
24h
Gelsolin-GFP
c
Control
24h
Average
Distance (μm)
d
5 mg ml-1
24h
Average
Distance (μm)

Figure S4

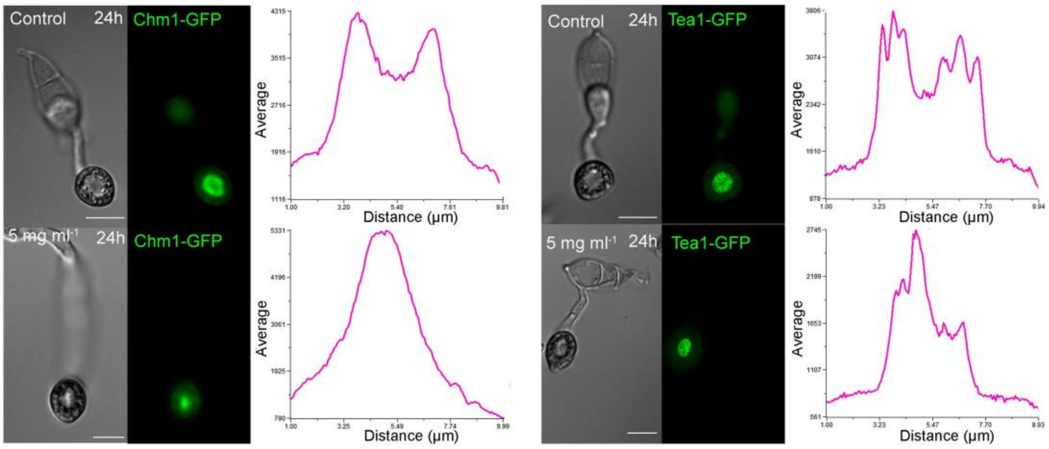
Control
24h
Chm1-GFP
5 mg ml-1
24h
Chm1-GFP
Average
Distance (µm)
Control
24h
Tea1-GFP
5 mg ml-1
24h
Tea1-GFP
Average
Distance (µm)

Figure S5

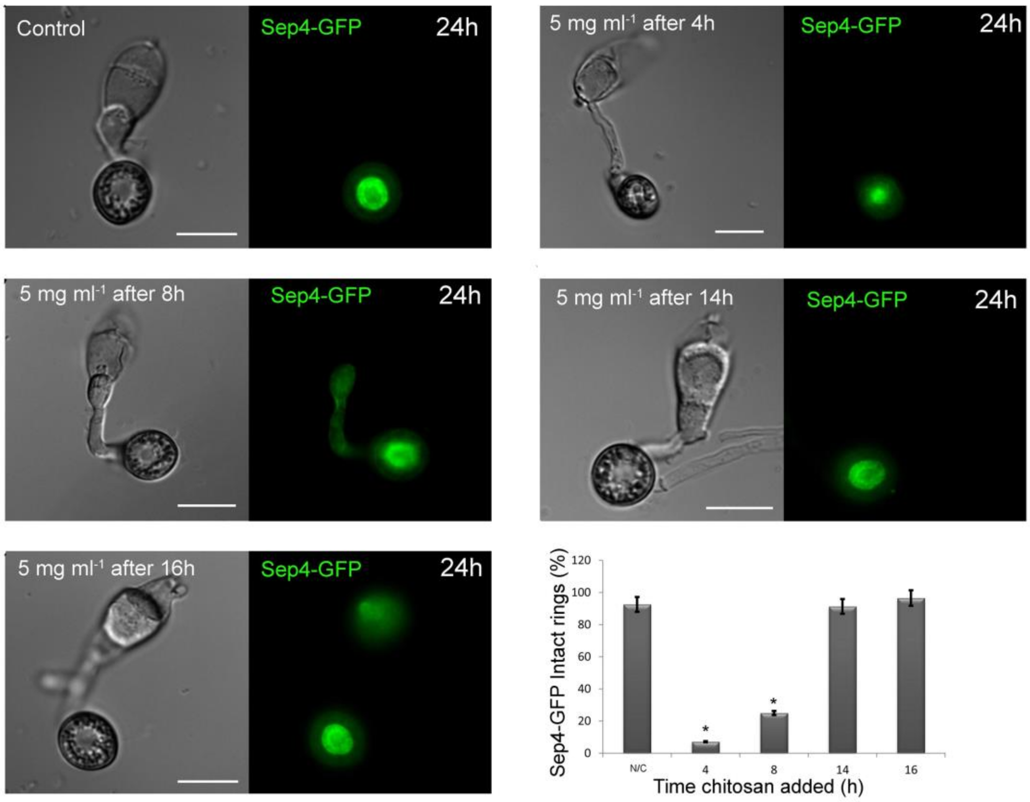
Control
Sep4-GFP
24h
5 mg ml-1 after 4h
Sep4-GFP
24h
5 mg ml-1 after 8h
Sep4-GFP
24h
5 mg ml-1 after 14h
Sep4-GFP
24h
5 mg ml-1 after 16h
Sep4-GFP
24h
Sep4-GFP Intact rings (%)
120
100
80
60
40
20
0
N/C
4
8
14
16
Time chitosan added (h)

Figure S6

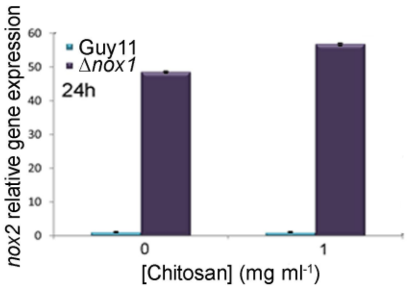
nox2 relative gene expression
60
50
40
30
20
10
0
Guy11
Δnox1
24h
0
1
[Chitosan] (mg ml-1)

Figure S7

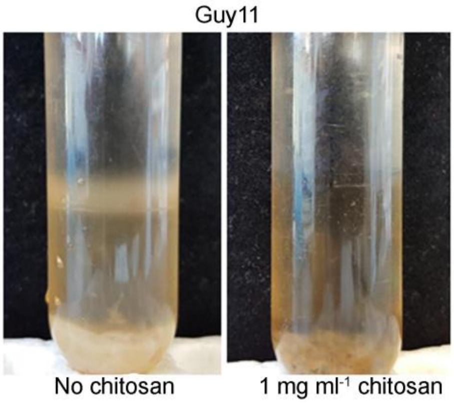


Figure S8
